# Retinoic acid signaling in mouse retina endothelial cells is required for early angiogenic growth

**DOI:** 10.1101/2022.05.09.491241

**Authors:** Christina N. Como, Cesar Cervantes, Brad Pawlikowski, Julie Siegenthaler

## Abstract

The development of the retinal vasculature is essential to maintain health of the tissue, but the developmental mechanisms are not completely understood. The aim of this study was to investigate the cell-autonomous role of retinoic acid signaling in endothelial cells during retina vascular development. Using a temporal and cell-specific mouse model to disrupt retinoic acid signaling in endothelial cells in the postnatal retina (*Pdgfb^icre/+^ dnRAR403^fl/fl^* mutants), we discovered that angiogenesis in the retina is significantly decreased with a reduction in retina vascularization, endothelial tip cell number and filipodia, and endothelial ‘crowding’ of stalk cells. Interestingly, by P15, the vasculature can overcome the early angiogenic defect and fully vascularized the retina. At P60, the vasculature is intact with no evidence of retina cell death or altered blood retinal barrier integrity. Further, we identified that the angiogenic defect seen in mutants at P6 correlates with decreased *Vegfr3* expression in endothelial cells. Collectively, our work identified a previously unappreciated function for endothelial retinoic acid signaling in early retinal angiogenesis.

## INTRODUCTION

Development of the blood vasculature that supplies the retina is a highly orchestrated process that is essential for normal vision. Neurovascular congruence is a process by which the nervous system co-develops with the blood supply, though the molecular details remains incompletely understood (1). The retina has a precisely layered neuronal architecture in which ganglion cells are in the inner retina, and light-detecting photoreceptors are in the outer retina (2). The retinal vasculature is divided into three layers, the primary, intermediate, and deep plexuses; the primary layer is adjacent to ganglion cells, and the deep vessels are next to photoreceptor nuclei. In addition to adjacent layering of neuronal and vascular elements, there is significant overlap in retinal neurogenesis and growth and remodeling of the vascular plexuses. This suggests that developmental signals important for neuronal development in the retina may also play a role in development of the retinal vasculature.

Initiation of retina angiogenesis is triggered by tissue hypoxia and is followed by upregulation and release of a proangiogenic factor, vascular endothelial growth factor (VEGF) primarily by astrocytes but also Müller cells, radial glial cells, retinal pigment epithelial cells, and photoreceptor cells(3). VEGF acts on endothelial cells within the retina to initiate and drive angiogenesis (4). Growth of new vessels depends on endothelial cells making vascular sprouts known as endothelial tip cells with long dynamic filipodia (5). Behind the tip cells are endothelial stalk cells that are highly proliferative and form the nascent vascular lumen (4,6). A balance of VEGF and Notch signaling pathways regulates the ratio of tip cell and stalk cells to permit progression of the primary vascular plexus (5,7). Growth of the secondary and tertiary vascular plexus depends on Notch and VEGF, but also myeloid cell derived Wnt ligands (8). Vessel growth in the primary plexus is followed by vascular remodeling, this is achieved by vessel ‘pruning’ whereby vessels are retracted, leaving behind an extracellular matrix ‘sleeve’, frequently visualized by antibody staining for collagen 4 (Collagen-4). The cellular and molecular mechanisms underlying vessel pruning in the retina are not as well understood but are known to play a role in pruning excessive blood vasculature due to low blood flow conditions (9,10).

Retinoic acid (RA) signaling is required for normal eye development and there is emerging evidence it plays a role in retinal vascular and blood-retinal barrier (BRB) development and maintenance. RA is a hormone generated locally within tissues via biosynthetic processing of Vitamin A by retinaldehyde dehydrogenases (RALDH1, 2, 3) and induces cell signaling via binding to nuclear hormone receptors, retinoic acid receptors (RAR) and retinoic X receptors (RXR) (11). RA is produced by multiple cells of the retina starting early in development, but not by retinal pigment epithelial cells (RPE) or cells of the choroid (12–14). RA signaling is essential for initial embryonic eye patterning and formation (15,16). In the postnatal mouse retina, RA is produced by multiple neural cells in the retina where it is required for proper eye development and maintenance (1). Studies in mouse and zebrafish using pharmacological inhibitors of RA synthesis or signaling support a role for RA in retinal blood vessel formation and BRB development (17,18). However, it is unclear from these studies if RA signaling is required in retinal endothelial cells or other retinal cell types to support vascular development. Animal models using targeted, cell-type specific perturbation of RA signaling is important to parse out the role of RA in retinal angiogenesis and BRB development.

To block RA signaling specifically in endothelial cells, we used endothelial cell CreERT mouse line (*Pdgfb^icre/+^*) to express dominate negative retinoic acid receptor (*dnRAR403^fl/fl^*) (19) in endothelial cells starting at early postnatal stages, prior to the start of retinal angiogenesis. Using retinal wholemounts, sections and endothelial cell isolation for gene expression profiling, we identify an early role for endothelial RA signaling in primary vascular plexus angiogenesis via regulation of tip cell number, potentially mediated by regulation of *Vegfr3*. Later vascular growth and BRB development are unaffected however persistent Collagen-4 sleeves in adult retinas suggests potential defects in vascular remodeling. Collectively our work supports a temporally restricted role for endothelial RA signaling in retinal vascular development.

## METHODS

### Animals

Mice used for experiments here were housed in specific-pathogen-free facilities approved by AALAC and were handled in accordance with protocols approved by the University of Colorado Anschutz Medical Campus IACUC committee. The following mouse lines were used in this study: *PdgfbiCre(20), dnRAR403-flox (21).* Mice were maintained on a mixed background and breeders (*Pdgfb^icre/+^, Pdgfb^icre/+^ dnRAR403^fl/fl^, dnRAR403^fl/+^*, and *dnRAR403^fl/fl^* adult males and females) used to generate experimental animals were generated via sibling crosses. All breeders were genotyped for the *Rd1* allele, this is absent from our background. To activate Cre-mediated recombinase activity, Tamoxifen (Sigma, St. Louis, MO, United States) was dissolved in corn oil (Sigma, St. Louis, MO, United States; 1mg/ml) and 20 ng was injected into pups at postnatal (P) day 1-3 to generate tamoxifen exposed *wildtype* (*WT*), *Pdgfb^icre/+^, dnRAR403^fl/fl^, and Pdgfb^icre/+^ dnRAR403^fl/fl^* genotypes. For all experiments, *wildtype* (*WT*) and *Pdgfb^icre/+^* samples were littermates whereas *dnRAR403^fl/fl^ and Pdgfb^icre/+^ dnRAR403^fl/fl^* samples were littermates.

### Whole Retina dissection and Immunohistochemistry

Mice were collected at P6, P15, and P60. For whole mount retina staining, eyes were dissected into cold PBS, retinas were removed, and fixed overnight in 4% paraformaldehyde at 4°C. Retinas were washed with 1x PBS and blocked in 3% bovine serum albumin (BSA) + 0.05% Triton-X overnight at 4°C. Retinas were stained with the following primary antibodies at a 1:100 dilution: Collagen-4 (BioRad, catalog # 2150-1470, Hercules, CA), ERG (Abcam, catalog # ab92513, Cambridge, UK), Iba1 (Cell Signaling, catalog # 17198, Danver, MA), desmin (Cell Signaling, catalog # 5332), or GFAP (Cell Signaling, 3670). Following incubation with primary antibody(s), sections were incubated with appropriate Alexafluor-conjugated secondary antibodies (Invitrogen, Carlsbad, CA, United States), Alexafluor 633-conjugated isolectin-B4 (Ib4; Invitrogen, Carlsbad, CA, United States), and DAPI (Invitrogen, Carlsbad, CA, United States). Immunofluorescent (IF) images were captured using a Zeiss (Thornwood, NY, United States) 900 LSM confocal microscope. Laser power and gain settings were always the same between control and mutant samples to accurately analyze expression of protein of interest.

### Cryosection

Whole eyeballs were collected, the cornea and lens were removed, and eyes were fixed overnight in 4% paraformaldehyde at 4°C. Retinas were washed with 1x PBS and cryoprotected with 20% sucrose in PBS overnight and subsequently frozen in OCT.

Tissue was cryosectioned in 12 μm increments and tissue sections used in analysis were matched for region using optic nerve. Sections were blocked in in 3% bovine serum albumin (BSA) + 0.05% Triton-X for 1 hour at room temperature and incubated with the following primary antibodies at a dilution of 1:100: Collagen-4, Fibrinogen (Abcam, catalog # ab34269), or PLVAP (BioRad, catalog # MCA2539GA). Following incubation with primary antibody(s), sections were incubated with appropriate Alexafluor-conjugated secondary antibodies (Invitrogen, Carlsbad, CA, United States), Alexafluor 633-conjugated isolectin-B4 (Ib4; Invitrogen, Carlsbad, CA, United States), and DAPI (Invitrogen, Carlsbad, CA, United States).

### Image Analysis

To quantify the amount of vascular expansion (microns), ImageJ was used to draw a line from the center of the optic nerve to the most radial point where blood vessels occupied the retina in WT (n=3), *Pdgfb^icre/+^* (n=3), *dnRAR403^fl/fl^* (n=6), and *Pdgfb^icre/+^* dnRAR403*^fl/fl^* (n=6) animals. To quantify the % vessel area, Angiotool (22) was used to quantify the amount of blood vessel area present in each image of the same magnification and made relative to WT animals in all controls and experimental: WT (n=5), *Pdgfb^icre/+^* (n=7), *dnRAR403*^fl/fl^ (n=10), and *Pdgfb^icre/+^ dnRAR403*^fl/fl^ animals (n=7). To quantify the number of Collagen 4 sleeves present, the number of collagen 4 sleeves (Collagen-4+, isolectinb4-) in each frame of equal magnification was quantified and tracked using ImageJ in WT (n=5), *Pdgfb^icre/+^* (n=3), *dnRAR403*^fl/fl^ (n=4), and *Pdgfb*^icre/+^ *dnRAR403*^fl/fl^ animals (n=3). The number of tip cells per frame were quantified using ImageJ to track all cells at the vascular front with extended filipodia, as well as the number of filipodia extended from each tip cell in WT (n=3), *Pdgfb*^icre/+^ (n=6), *dnRAR403*^fl/fl^ (n=4), and *Pdgfb^icre/+^ dnRAR403*^fl/fl^ (n=6) animals. The filipodia length was measured using ImageJ by drawing a line from the base of the tip cell to the most radial point of the filipodia in *dnRAR403*^fl/fl^ (n=4), and *Pdgfb*^icre/+^ *dnRAR403*^fl/fl^ (n=4) animals. To quantify the number of ERG overlapping nuclei, ImageJ was used to track the number of nuclei that were close enough in proximity to overlap in WT (n=3), *Pdgfb*^icre/+^ (n=3), *dnRAR403*^fl/fl^ (n=3), and *Pdgfb*^icre/+^ *dnRAR403*^fl/fl^ (n=3) animals. The number of proliferating ERG cells was quantified using ImageJ to track and determine ERG+EDU+ cells in *dnRAR403*^fl/fl^ (n=3), and *Pdgfb*^icre/+^ *dnRAR403*^fl/fl^ (n=3) animals. The number of microglia was quantified by counting the number of Iba1+ cells in frames of equal magnifications in *dnRAR403*^fl/fl^ (n=5), and *Pdgfb*^icre/+^ *dnRAR403*^fl/fl^ (n=4) animals. The % pericyte blood vessel coverage was quantified using ImageJ by taking (pericyte area/total blood vessel area) and multiplying it by 100 in *dnRAR403*^fl/fl^ (n=3), and *Pdgfb*^icre/+^ *dnRAR403*^fl/fl^ (n=6) animals. To quantify the % astrocyte blood vessel coverage, ImageJ was used to determine the amount of (astrocyte-vessel area overlap/total astrocyte area) in *dnRAR403*^fl/fl^ (n=5) and *Pdgfbi*^cre/+^ *dnRAR403*^fl/fl^ (n=5) animals. To determine the width of neuron layers within retina sections, ImageJ was used to quantify the width of each neuronal layer in microns in in *dnRAR403*^fl/fl^ (n=3), and *Pdgfb*^icre/+^ *dnRAR403*^fl/fl^ (n=3) animals.

### Endothelial Cell Enrichment

To enrich endothelial cells from whole retinas, retinas were dissected on ice, cut into fine pieces using forceps, and placed in DMEM serum free media with 1 mg/mL collagenase and 10 ug/mL DNAse for 15 minutes at 37°C. Retinas were then broken up using a 1000 mL pipette. An equivalent volume of DMEM + 10% fetal bovine serum was added to deactivate the enzymes. Isolation of endothelial cells was then completed using the MACS cell isolation protocol magnetic CD31 beads (Miltenyi, Bergisch Gladbach, Germany).

### RT-qPCR

RNA from isolated endothelial cells was obtained using the Qiagen RNAeasy kit (Hilden, Germany). cDNA was then synthesized using iScript cDNA synthesis kit (BioRad, Hercules, CA, United States) and RT-qPCR was performed to ensure endothelial cell enrichment using endothelial specific markers (*Cldn5, Sox17, Tie2, Cdh5*) and that other markers from other cells were depleted (Map2, GFAP) relative to cDNA isolated from whole retina. Analysis of differential signaling was quantified using primers to *Pdgfb, Cdh5* (VE-cadherin), *Hes1, Hey1, Notch1, Dll4, Lef1, Axin2, Vegfr2*, and *Vegfr3* (Integrated DNA Technologies, Coralville, Iowa). *Actb* transcript levels were also assessed and used to normalize expression levels. Delta-delta Ct analysis was performed and fold change over control is reported. All sequences of primers used are provided in Table 1.

**Table 1.**
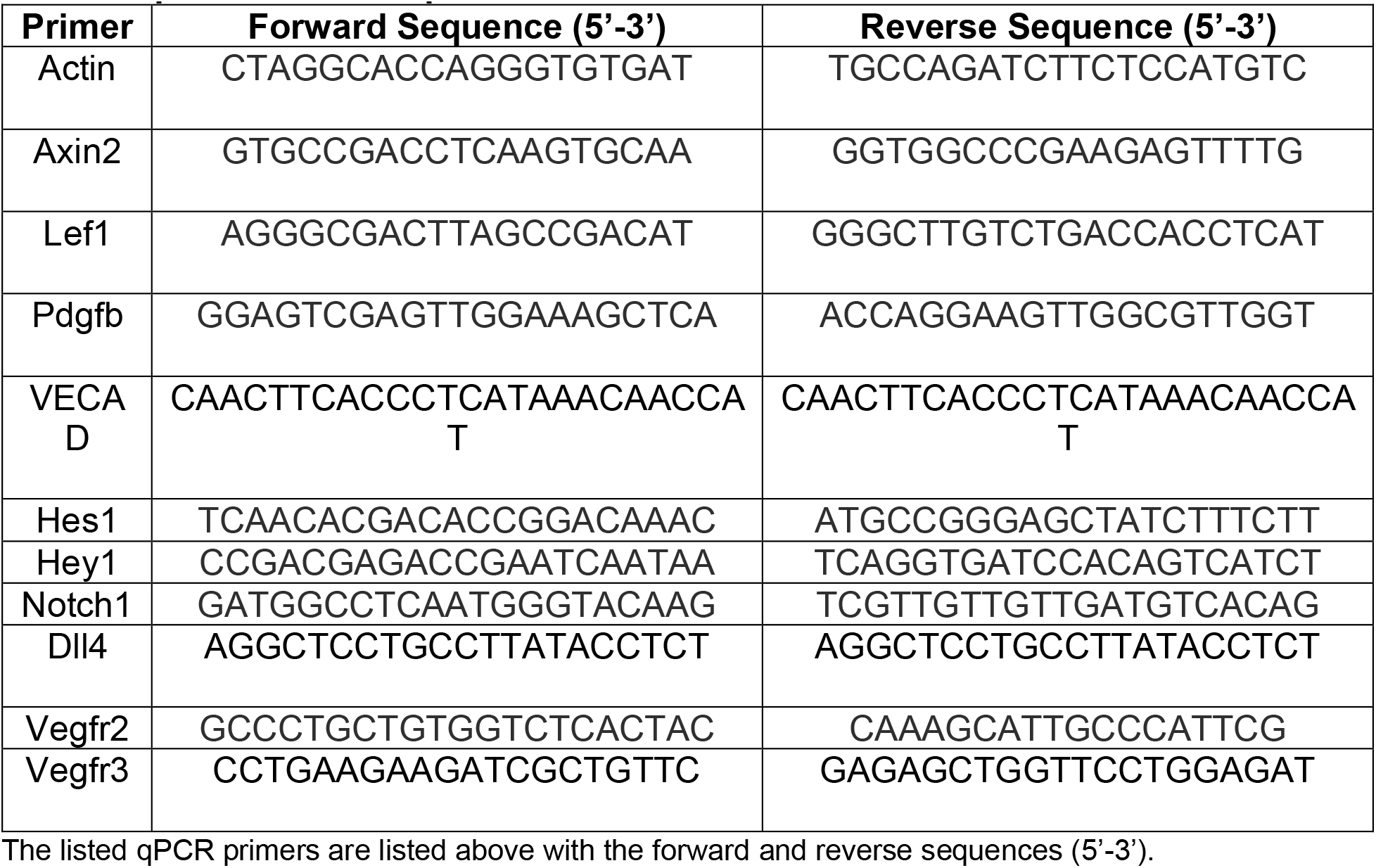
qPCR Primer Sequences.

### Statistics

To detect statistically significant differences in mean values of controls and mutant genotypes (vascular expansion, vessel area, collagen 4 sleeves, number of tip cells, number of stacked ERG cells, neuron layer length) a non-parametric Kruskal Wallis with a Dunn’s multiple comparisons test was used to detect statistically significant differences between treatment conditions. For comparison of two values (# of filipodia, proliferating ERG cells, # of microglia, pericyte vessel coverage, astrocyte vessel coverage), a Mann-Whitney was used. For comparison of endothelial cell volume and filipodia length, and RT-qPCR gene expression, a two-way ANOVA was used. *P*-values less than 0.05 were considered statistically significant in these studies with specific *p*-values reported on all graphs. The standard deviation (SD) is reported on all graphs.

## RESULTS

### At postnatal (P) day 6, *Pdgfb^icre/+^ dnRAR403^fl/fl^* retinas have defective angiogenesis

Postnatal (P) day 1-3 mice were injected with tamoxifen and analyzed at P6. Compared to controls (wildtype, *Pdgfb^icre/+^* and *dnRAR403^fl/fl^*, mice), *Pdgfb^icre/+^dnRAR403^fl/fl^* retinas were not as vascularized (Figure 1A; DAPI, blue; collagen 4, red; PECAM, white). Quantification of vascular distance (length from the optic nerve to the vascular front) revealed that compared to controls, *Pdgfb^icre/+^ dnRAR403^fl/fl^* retinas had a significantly reduced vascular distance (Figure 1B, 488.6 ± 51.95, 461.2 ± 53.24, 416.2 ± 56.89, 318.8 ± 29.44, respectively; mean vascular distance in microns ± SD). Quantification of the blood vessel area using Angiotool revealed that compared to controls, *Pdgfb^icre/+^ dnRAR403^fl/fl^* retinas had significantly less blood vessel area within the primary plexus (Figure 1C, 1.0 ± 0.17, 0.94 ± 0.21,0.95 ± 0.08, 0.62 ± 0.06, respectively; mean % blood vessel area relative to WT ± SD). Analysis of the presence of empty basement membrane vessels were quantified by looking at the number of Collagen-4 sleeves by staining for lectin and Collagen-4 to label blood vessels. The number of collagen-4 sleeves were quantified as collagen-4 positive and lectin negative (Figure 1D, yellow arrow). Compared to controls, *Pdgfb^icre/+^ dnRAR403^fl/fl^* retinas had significantly reduced collagen-4 sleeves (Figure 1E, 37.0 ± 3.08, 36.33 ± 4.73, 40.58 ± 6.82, 10.53 ± 0.65; mean # of collagen-4 sleeves per frame ± SD). Analysis of individual endothelial cell size was visualized using claudin-5 staining, a tight junction protein that marks where two endothelial cells are joined and can be used to approximate the outside border of individual endothelial cells (Figure 1F, yellow dashed line). Control *dnRAR403^fl/fl^* mice and mutant *Pdgfb^icre/+^ dnRAR403^fl/fl^* endothelial cells did not have a significant difference in individual endothelial cell size (Figure 1G). Overall, this data shows there is not a Cre or dnRAR alone genetic phenotype in control animals, and that the angiogenic deficits seen at P6 is due to the disruption of RA signaling in endothelial cells. The disruption of RA signaling in endothelial cells causes a vascular expansion defect, characterized by reduced blood vessel expansion and blood vessel area but normal endothelial cell size. Decreased collagen-4 sleeves (a read-out of vascular retraction and remodeling) may result from a general reduction in vascular plexus density in mutant *Pdgfb^icre/+^dnRAR403^fl/fl^* samples.

**Figure 1.**
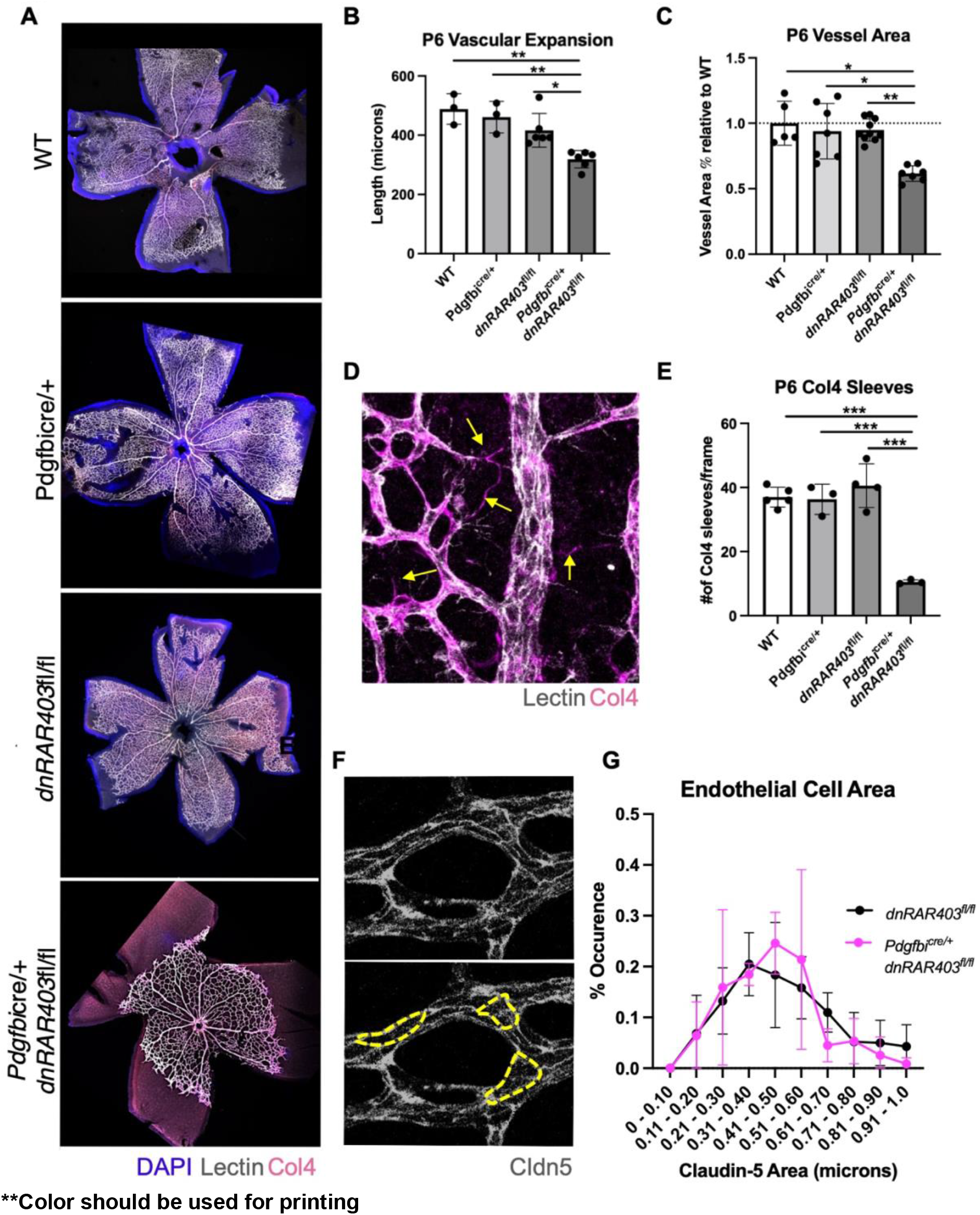
Postnatal angiogenesis is significantly reduced in *Pdgfb^icre/+^ dnRAR403^fl/fl^* mice. Postnatal (P) day 1-3 mice were injected with tamoxifen and analyzed at P6. Compared to wildtype (WT), *Pdgfb^icre/+^*, and *dnRAR403^fl/fl^* mice, *Pdgfb^icre/+^ dnRAR403^fl/fl^* retinas (A; DAPI, blue; collagen 4, magenta; PECAM, white) had significantly reduced vascular expansion (B). Analysis of the primary plexus blood vessel area revealed that compared to controls, *Pdgfb^icre/+^ dnRAR403^fl/fl^* retinas had significantly less blood vessel area (C). Analysis of the presence of collagen-4 (Collagen-4) sleeves (D, magenta) revealed that compared to controls, *Pdgfbicre/+ dnRAR403^fl/fl^* retinas have significantly decreased Collagen-4 sleeves at P6 (E). Endothelial cells were stained with claudin-5 (F, white) and the outside of each endothelial cell was outlined (yellow dashes) to quantify endothelial cell area. There was no significant difference in endothelial cell volume between *dnRAR403^fl/fl^* controls and *Pdgfb^icre/+^ dnRAR403^fl/fl^* mutants. (p <0.05, *; p<0.01, **; p<0.001, ***).

### At P6, *Pdgfb^icre/+^ dnRAR403^fl/fl^* have a disruption in tip cell number and filipodia

We next tested if diminished vascular growth in mutant *Pdgfb^icre/+^ dnRAR403^fl/fl^* was due to alterations in angiogenesis features at the primary plexus vascular front, specifically tip cells number and tip cell filipodia extensions. The number of tip cells at the vascular front was visualized in retinal whole mounts of controls (wildtype, *Pdgfb^icre/+^,* and *dnRAR403^fl/fl^* mice), and experimental mice (*Pdgfb^icre/+^ dnRAR403^fl/fl^* mice) (Figure 2A; lectin, white; yellow asterisk indicates tip cell). Quantification of the number of tip cells at the vascular front revealed that compared to controls, *Pdgfb^icre/+^ dnRAR403^fl/fl^* retinas had significantly reduced tip cells (Figure 2B; 12.44 ± 1.08, 11.83 ± 1.69, 13.65 ± 2.20, 6.40 ± 0.97; mean # of tip cells/frame ±SD). The number of filipodia extended from each tip cell was quantified and compared to *dnRAR403^fl/fl^* retinas. *Pdgfb^icre/+^ dnRAR403^fl/fl^* tip cells had significantly decreased filipodia (Figure 2C, 4.00 ± 0.28, 2.05 ± 0.80; mean # of filipodia/tip cell ± SD). We measured filipodia length as a gross proxy for filipodia structure (Figure 2D, cyan dotted lines) and the quantification revealed that compared to *dnRAR403^fl/fl^* retinas, *Pdgfb^icre/+^ dnRAR403^fl/fl^* retinas did not have a significant difference in filipodia length measured between 5-70 microns (Figure 2E). This data suggests that RA signaling is required to generate tip cells and their respective filipodia, but that there is not an overt defect in the filipodia structure.

**Figure 2.**
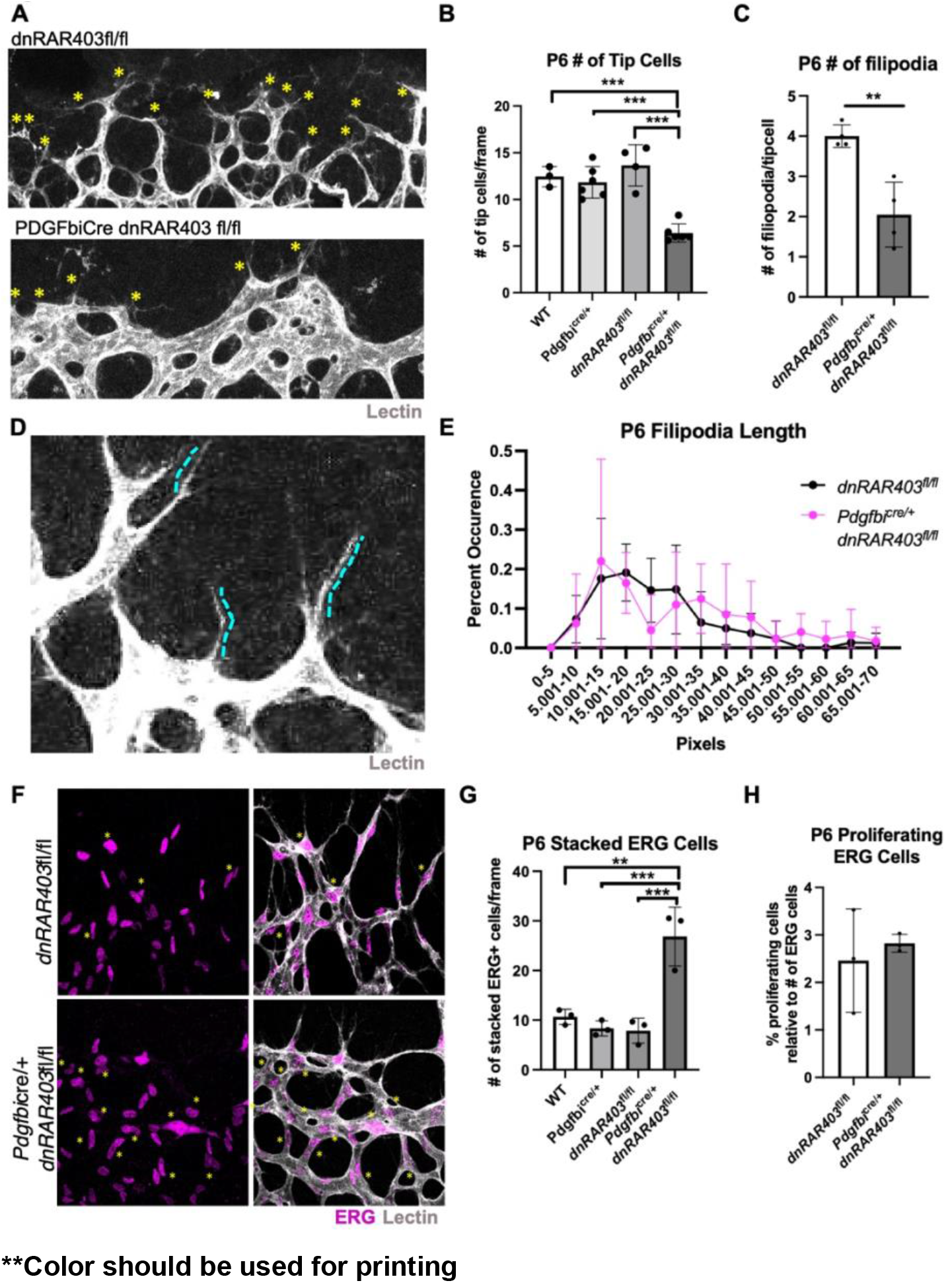
The loss of endothelial retinoic acid signaling reduces tip cell and filopodia number at the vascular front. Quantification of the number of tip cells at the vascular front (A, yellow asterisks) revealed that compared to controls, *Pdgfb^icre/+^ dnRAR403^fl/fl^* retinas had significantly reduced tip cells (B) and the number of filipodia extended from each tip cell (C). Quantification of the length of filipodia (D) between controls and *Pdgfb^icre/+^ dnRAR403^fl/fl^* retinas revealed no significant difference (E). Analysis of the number of stacked endothelial cells (F, yellow asterisk) stained with ERG (magenta) and vessels stained with IB4 (white) revealed that compared to controls, *Pdgfb^icre/+^ dnRAR403^fl/fl^* retinas have significantly increased overlapped ERG nuclei. The number of proliferating endothelial cells, as labeled with EdU and ERG, were not significantly different between control and *Pdgfb^icre/+^ dnRAR403^fl/fl^* retinas. (p <0.05, *; p<0.01, **; p<0.001, ***).

### *Pdgfb^icre/+^ dnRAR403^fl/fl^* mutants at P6 have ‘stacked’ endothelial cells but proliferate normally

To determine if endothelial RA signaling has a role in growth of stalk cells near the vascular front, we quantified the organization and proliferation of stalk cells. To quantify endothelial cell organization, we scored the number of ERG positive nuclei that were overlapping with another ERG+ nuclei in a defined area (Figure 2F, yellow asterisk; lectin, white), referred to as ‘stacked’ endothelial cells. Quantification of the number of stacked endothelial cells at the vascular front revealed that compared to controls (wildtype, *Pdgfb^icre/+^*, and *dnRAR403^fl/fl^* mice), *Pdgfb^icre/+^ dnRAR403^fl/fl^* retinas had significantly increased stacked ERG+ endothelial cells (Figure 2G, 10.67 ± 1.53, 8.33 ± 1.53, 7.89 ± 2.52, 26.83 ± 5.92; mean number of ERG+ stacked endothelial cells ± SD, respectively). The number of proliferating ERG+ endothelial cells were labeled with EdU and compared to dnRAR^fl/fl^ retinas, *Pdgfb^icre/+^ dnRAR403^fl/fl^* retinas did not have a significant difference in the number of proliferating endothelial cells (Figure 2H; 2.46 ± 1.09, 2.83 ± 0.18, respectively; mean # of proliferating ERG cells/frame ± SD). Overall, this data suggests that endothelial proliferation is not altered in endothelial RA signaling mutants but there is altered endothelial cell organization near the vascular front.

### Microglia, pericytes and astrocytes are not impacted by the loss of retinoic acid signaling in endothelial cells

To determine if other cell types contribute to the angiogenic phenotype seen in endothelial RA signaling mutants, we investigated microglia, pericytes, and astrocytes. Compared to *dnRAR403^fl/fl^* control retinas, *Pdgfb^icre/+^ dnRAR403^fl/fl^* retinas did not have a significantly different number of microglia (Figure 3A, 28.40 ± 4.16, 30.75 ± 2.99, mean # of microglia/frame ± SD) or qualitatively in morphology (Figure 3B; Iba1, cyan; lectin, magenta). The percent pericyte-blood vessel coverage was not significantly different between *dnRAR403^fl/fl^* control retinas or *Pdgfb^icre/+^ dnRAR403^fl/fl^* retinas (Figure 3C, 14.34 ± 3.62, 16.93 ± 5.90; mean % (pericyte area/blood vessel area) ± SD). There was not a qualitatively observed morphological difference between pericytes in controls and mutants (Figure 3D; desmin, cyan; lectin, magenta). The area of astrocyte blood vessel coverage was not significantly different between controls or mutants (Figure 3E, 37.11 ± 5.52, 29.34 ± 8.16, mean (astrocyte blood vessel area/ blood vessel area) ± SD) or in morphology (Figure 3F; lectin, magenta; GFAP, cyan). Thus, perturbation of RA signaling in endothelial cells does not overtly alter the morphology or blood vessel interaction of microglia, pericytes, or astrocytes.

**Figure 3.**
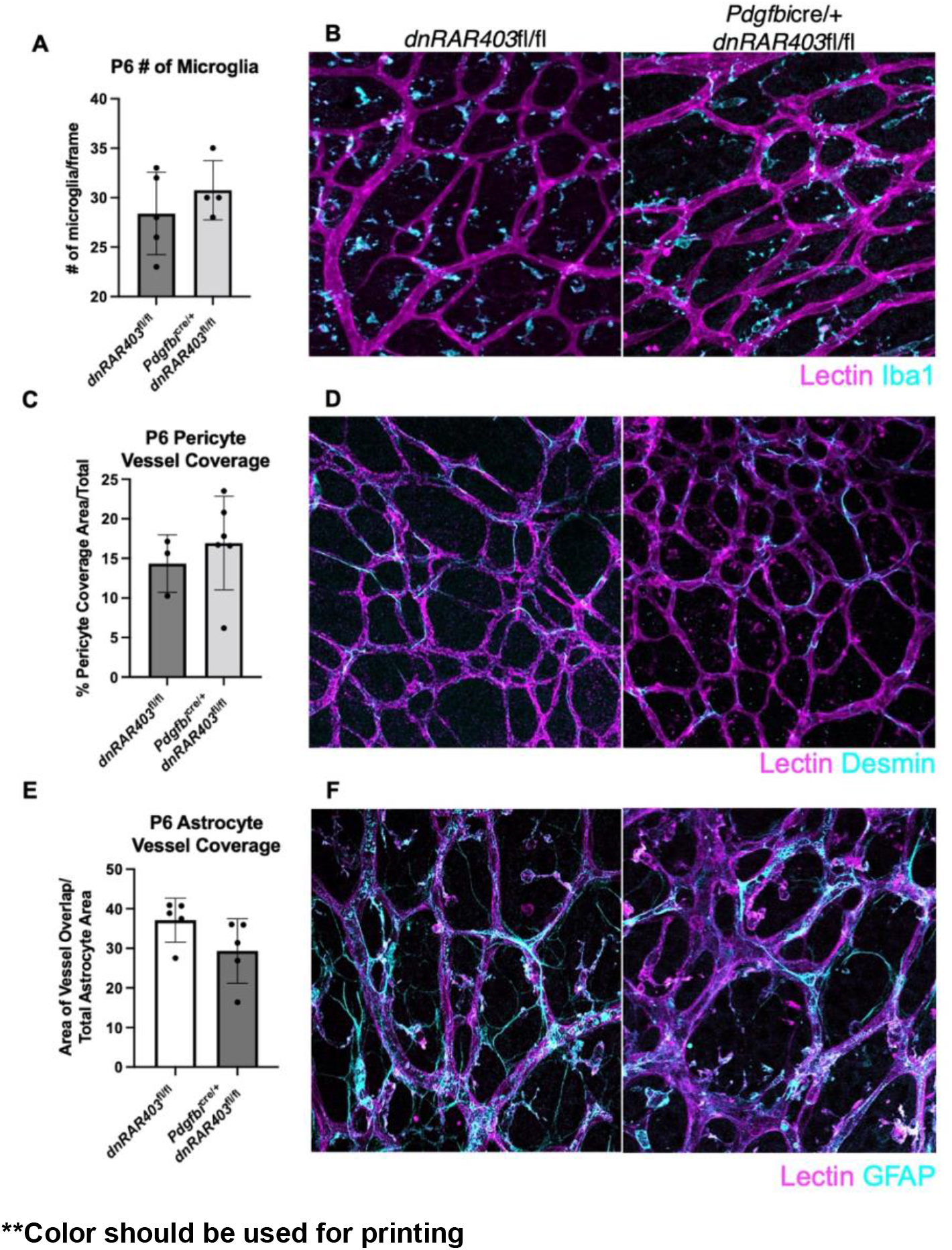
Microglia, pericytes and astrocytes are not overtly altered by the loss of retinoic acid signaling in endothelial cells. Like *Pdgfb^icre/+^* control mice, *Pdgfb^icre/+^ dnRAR403^fl/fl^* retinas display no quantitative difference in the number of microglia present (A) or the morphology (B, lectin, magenta; Iba1, cyan). The area of pericyte coverage of blood vessels was not significantly different between controls and Pdgfbicre/+ dnRAR^fl/fl^ retinas (C) or in morphology (D, lectin, magenta; desmin, cyan). The area of astrocyte vessel coverage was not significantly different between controls and Pdgfbicre/+ retinas (E) or in morphology (F, lectin, magenta; GFAP, cyan). (p <0.05, *; p<0.01, **; p<0.001, ***).

### Angiogenesis is recovered by P15 in *Pdgfb^icre/+^ dnRAR403^fl/fl^* retinas, but there are elevated number of collagen 4 sleeves

By P15, controls and *Pdgfb^icre/+^ dnRAR403^fl/fl^* retinas are fully vascularized (Figure 4A; DAPI, blue; lectin, white; Collagen-4, magenta) and had no significant differences in blood vessel area in the primary, intermediate, or deep plexus (Figure 4B; 1.00 ± 0.25 versus 1.36 ± 0.44, 1.00 ± 0.21 versus 1.07. ± 0.02, 1.00 ± 0.19 versus 1.04 ± 0.27, respectively; mean blood vessel area relative to control ± SD). Analysis of collagen 4 sleeves at P15 revealed that compared to *dnRAR403^fl/fl^* controls, *Pdgfb^icre/+^ dnRAR403^fl/fl^* retinas had significantly more sleeves in the primary plexus, no difference in the intermediate plexus, and significantly decreased sleeves in the deep plexus (Figure 4C; 4.17 ± 0.76 versus 30.00 ± 6.34; 5. 50 ± 2.78 versus 0.88 ± 0.75; 70.33 ± 28.30 versus 20.38 ± 7.17, respectively; mean Collagen-4 sleeves ± SD). At P60, compared to controls (wildtype, *Pdgfb^icre/+^, and dnRAR403^fl/fl^* mice), *Pdgfb^icre/+^dnRAR403^fl/fl^* retinas did not have a significantly different blood vessel area in the primary, intermediate, or deep plexus (Figure 4D). Compared to controls, *Pdgfb^icre/+^ dnRAR403^fl/fl^* retinas had significantly increased Collagen-4 sleeves in the primary plexus, but no significant difference in the intermediate or deep plexus (Figure 4E). Overall, this data suggests that there is not a long term Cre or genetic phenotype in our mice and that the deletion of retinoic acid signaling in endothelial cells does not have a persistent angiogenic phenotype at later time points. However, there are residual Collagen-4 sleeves in the primary plexus of adult mice.

**Figure 4.**
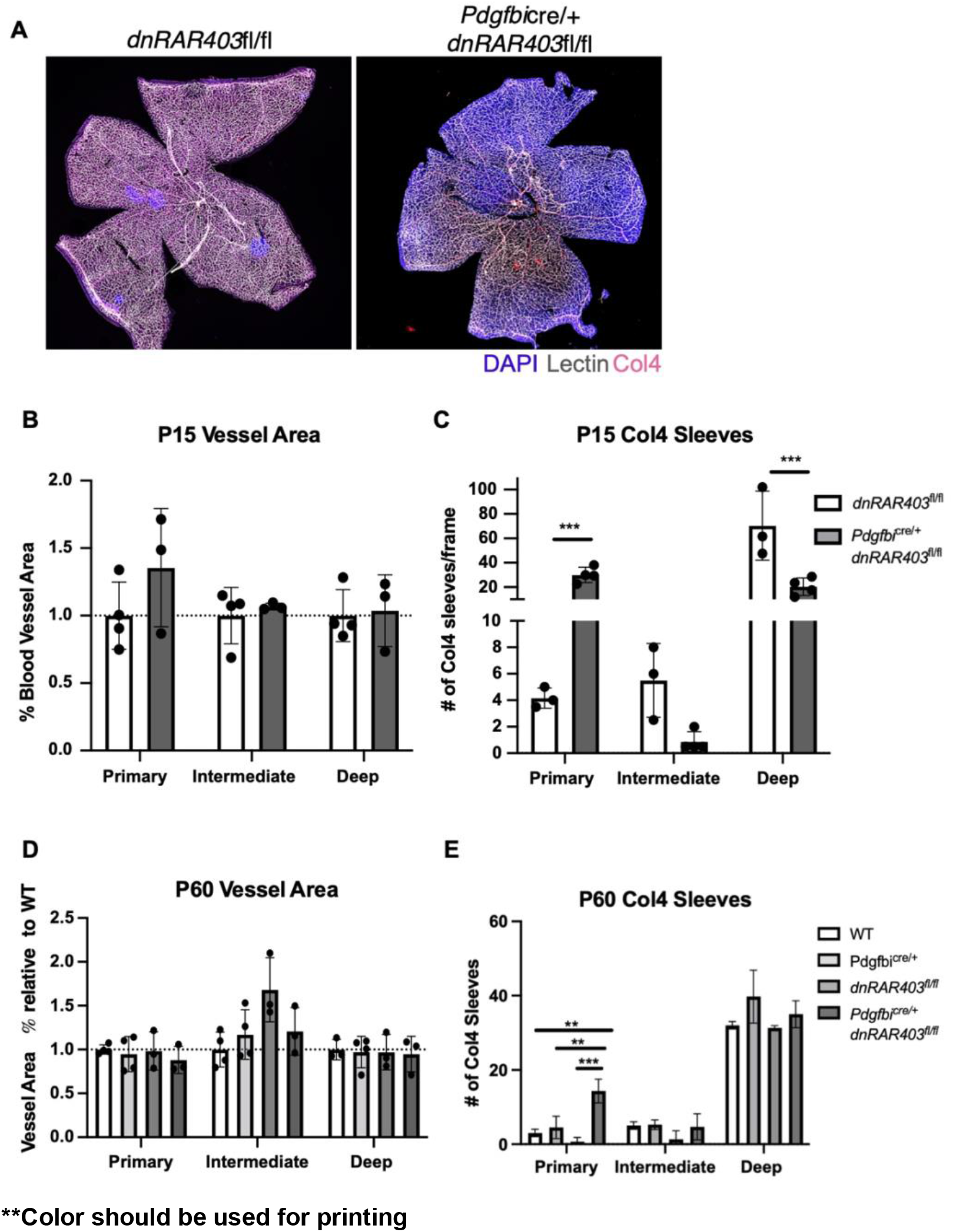
Angiogenesis recovers in *Pdgfb^icre/+^ dnRAR403^fl/fl^* retinas but there are defects in collagen 4 sleeves. Like controls, *Pdgfb^icre/+^ dnRAR403^fl/fl^* retinas are fully vascularized (A) with no significant difference in vessel area (B). Analysis of Collagen-4 sleeves at P15 revealed that there were significantly more Collagen-4 sleeves in the primary plexus, no difference in the intermediate plexus, and significantly decreased sleeves in the deep plexus (C). At P60, the blood vessel area was not significantly different from controls in the primary, intermediate, or deep plexus (D). Compared to controls, *Pdgfb^icre/+^ dnRAR403^fl/fl^* retinas had significantly increased Collagen-4 sleeves in the primary plexus, but no significant difference in the intermediate or deep plexus (E).

### Disruption of endothelial cell retinoic acid signaling does not affect BRB integrity or the number of neurons in adult mice

To determine if the deletion of RA signaling affects the tissue integrity retina long term or BRB integrity, postnatal (P) day 1-3 mice were injected with tamoxifen and analyzed at P60. Compared to controls, *Pdgfb^icre/+^ dnRAR403^fl/fl^* retinas did not have any obvious neuronal degeneration in any of the three neuronal layers (photoreceptors [PR], inner nuclear layer [INL], outer nuclear layer [ONL], Figure 5A; insets A’ and A’’). Analysis of neuronal cell layer thickness revealed that compared to controls, *Pdgfb^icre/+^dnRAR403^fl/fl^* retinas were not significantly different (Figure 5B). To determine if there is a disruption in blood-retinal barrier integrity, controls and mutants were stained with fibrinogen, a bloodborne protein that is normally excluded from CNS by an intact BBB and BRB (23,24). Controls (Figure 5C, left) and *Pdgfb^icre/+^dnRAR403^fl/fl^* retinas had no obvious fibrinogen signal in the retina (Figure 5C, right; DAPI, blue; lectin, magenta; fibrinogen, yellow). Additionally, there was no detectable endothelial PLVAP signal (Figure 5D, DAPI, blue; lectin, magenta; PLVAP, yellow), a pore protein that is not expressed in non-fenestrated, BBB and BRB endothelial cells, except for in PLVAP+ ciliary bodies (Figure 5E, yellow).

**Figure 5.**
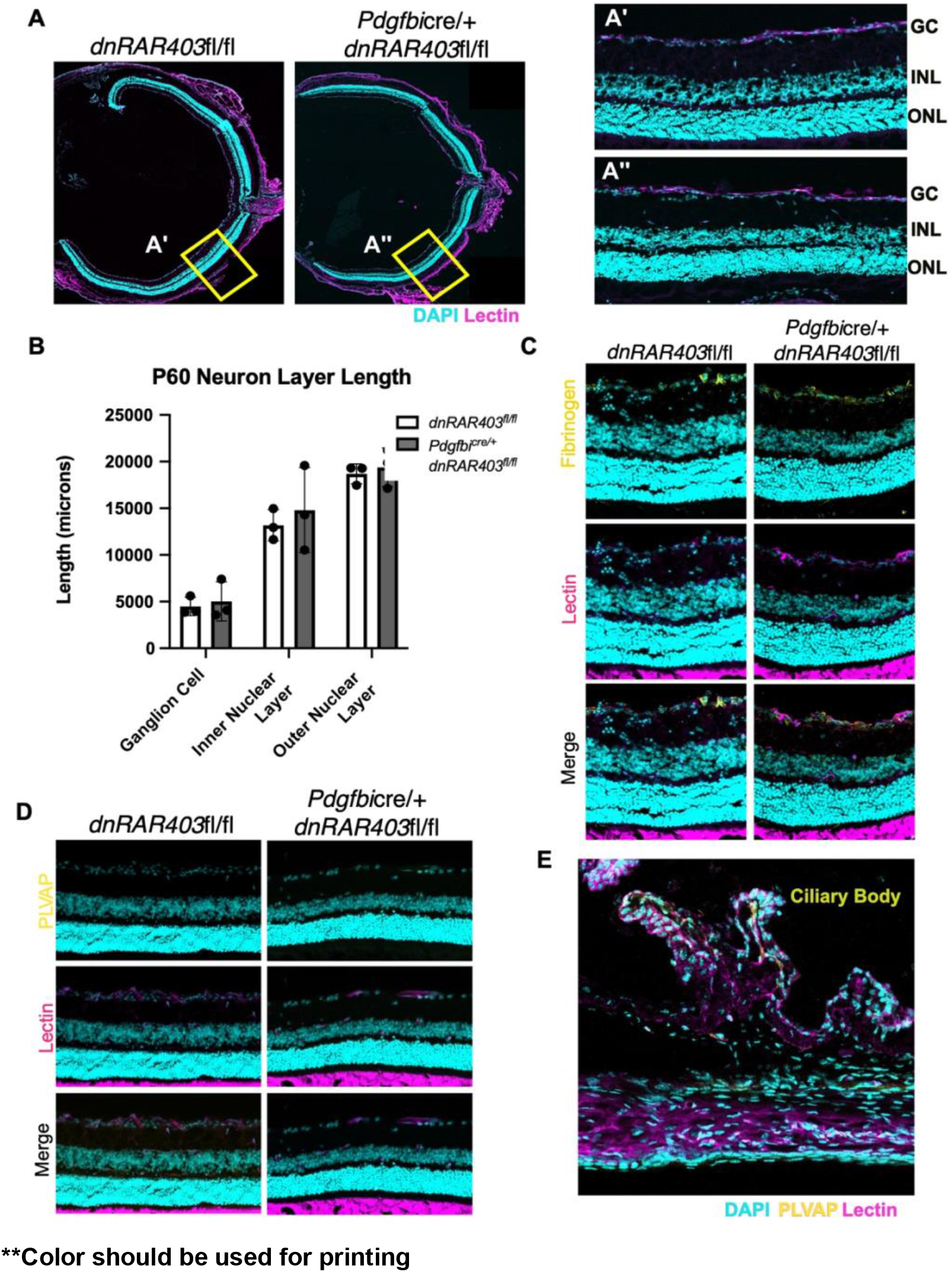
Disruption of endothelial cell retinoic acid signaling does not affect blood-retinal barrier integrity or the number of neurons in adult mice. Postnatal (P) day 1-3 mice were injected with tamoxifen and analyzed at P60. Compared to controls (A and A’), *Pdgfb^icre/+^ dnRAR403^fl/fl^* retinas (A and A”) have similar thickness in neuron layer ganglion cells, inner nuclear layer, or outer nuclear layer (B). BRB integrity was not overtly altered as there was no increase in fibrinogen (yellow) staining in the retina (C) or PLVAP in retinal vasculature (yellow) (D). As a positive control staining of PLVAP in same section ciliary bodies stained positive for PLVAP (yellow) (E). (p <0.05, *; p<0.01, **; p<0.001, ***).

### The loss of retinoic acid signaling in endothelial cells decreases *Vegfr3* but has no effect on *Vegfr2* or Notch pathway genes

To identify potential angiogenic pathways dysregulated in *Pdgfb^icre/+^ dnRAR403^fl/fl^* mutant retinas, we used PECAM-coated magnetic beads to enrich for endothelial cells, isolated RNA and used quantitative PCR for measure expression of Notch, Wnt and VEGF pathway related genes (Figure 6A, BioRender). To test for isolated endothelial cells from control *dnRAR403^fl/fl^* and mutant *Pdgfb^icre/+^ dnRAR403^fl/fl^* retinas were analyzed by RT-qPCR at P6. Relative to control expression levels, *Pdgfb^icre/+^ dnRAR403^fl/fl^* had no significant differences in *Pdgfb* (1.19 ± 0.72 versus 2.01 ± 0.45), *Hes1* (1.078 ± 0.48 versus 0.85 ± 0.1), *Hey1* (1.03 ± 0.26 versus 1.20 ± 0.20), *Notch1* (1.02 ± 0.24 versus 1.20 ± 0.43), *Dll4* (1.00 ± 0.10 versus 1.27 ±0.44), *Axin2* (1.02 ± 0.25 versus 1.41 ± 0.24), and *Vegfr2* (1.02 ± 0.22 versus 0.71 ± 0.20) and a significant difference in *Cdh5* (1.01 ± 0.72 versus 0.47 ± 0.13), *Lef1* (1.03 ± 0.26 versus 0.42 ± 0.10), and *Vegfr3* (1.02 ±0.22 versus 0.21 ± 0.02) (Figure 6B).

**Figure 6.**
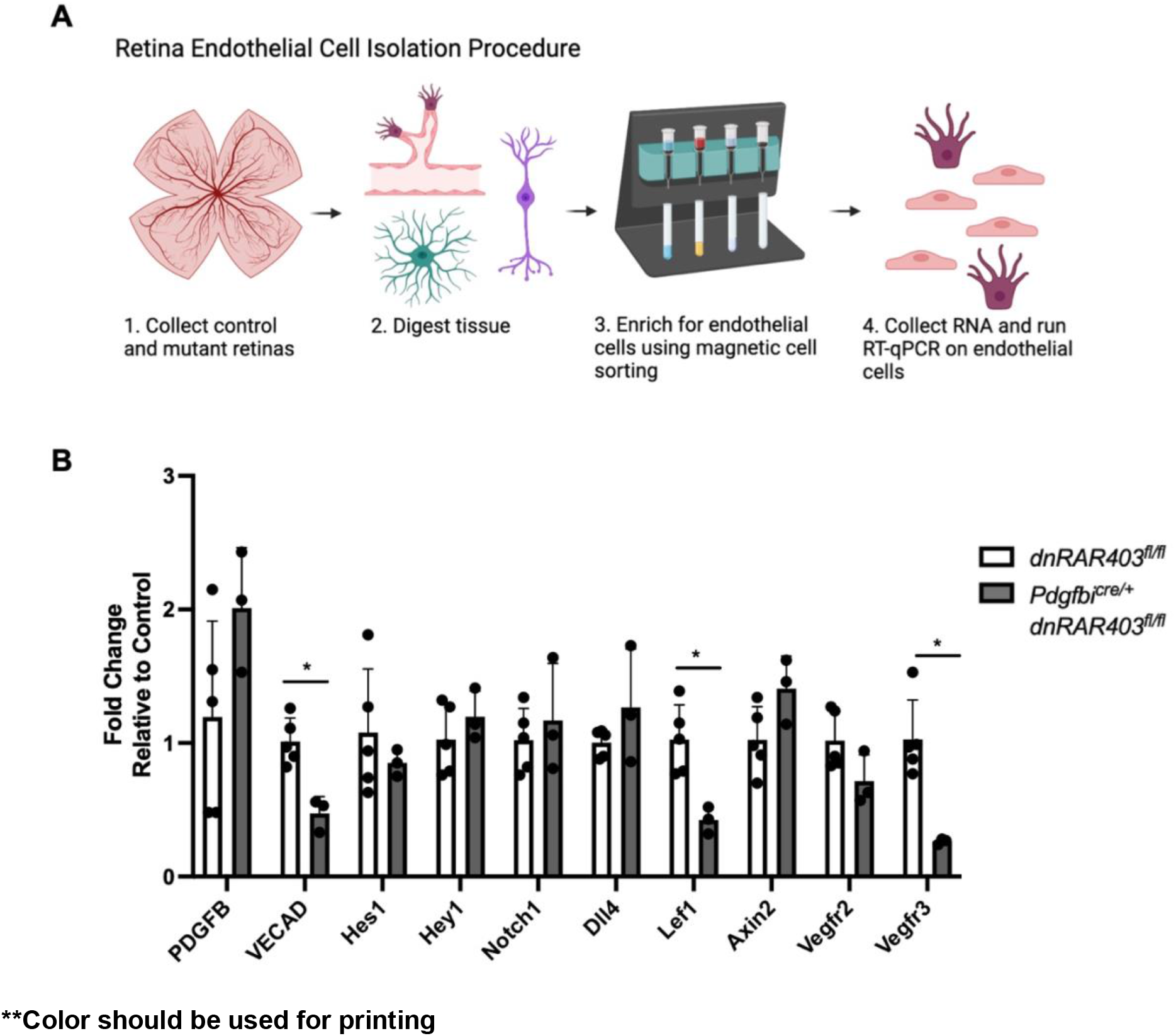
Gene expression profiling of retinal endothelial cells from control and *Pdgfb^icre/+^dnRAR403^fl/fl^*. (A) Endothelial cells were isolated from *dnRAR403^fl/fl^* and *Pdgfb^icre/+^ dnRAR403^fl/fl^* retinas and the RNA expression levels were quantified using RT-qPCR. Retinas were digested and endothelial cells were enriched using CD31 magnetic labeling. (B) Relative to controls, *Pdgfb^icre/+^ dnRAR403^fl/fl^* endothelial cells had significantly decreased levels of *Cdh5, Lef1*, and *Vegfr3*. There was no significant difference between controls and mutants of *Pdgfb, Hes1, Hey1, Notch1, Dll4, Axin2, or Vegfr2*. (p <0.05, *; p<0.01, **; p<0.001, ***).

## DISCUSSION

Here we describe a previously unknown function for endothelial RA signaling in the initial stages of vascular development. We show endothelial cell RA signaling promotes angiogenesis and primary plexus vascular expansion via regulation of tip cell number. Interestingly, development of the secondary and tertiary plexuses as well as BRB development does not rely on endothelial RA signaling. Further, we found that endothelial RA signaling may be upstream of endothelial VEGFR3, a VEGF receptor previously shown to modulate developmental angiogenesis. This reduction in angiogenic growth was not seen in wildtype, *Pdgfbi*^cre/+^ or *dnRAR403*^fl/fl^ control retinas. This is important as *Pdgfbi*^cre/+^ pups exposed to tamoxifen have slightly but significantly impaired development of the primary retinal vascular plexus (25). Differences in strain background may account for absence of phenotype in *Pdgfbi*^cre/+^ in our studies however their work underscores the importance of Cre and flox only controls for retinal angiogenesis studies in mice.

Our analysis of the angiogenic front and endothelial gene expression indicates that endothelial RA signaling regulates endothelial tip cells formation. Endothelial tip cells with filapodial sprouts “sense” the surrounding environment, responding to angiogenic cues such as VEGF that stimulate vascular expansion. Immediately behind the tip cells, are the trailing stalk cells that proliferate to help advance the vasculature toward the edge of the retina (26). We discovered that compared to controls (WT, *Pdgfbi*^cre/+^, and *dnRAR403*^fl/fl^), *Pdgfbi*^cre/+^ *dnRAR403*^fl/fl^ mutants have significantly decreased numbers of endothelial tip cells, as well as the number of filipodia per tip cell. However, the filopodial branches that were extended were not significantly different in length. Collectively, these data indicate that RA signaling plays an important role in tip cell selection. Interesting, we found no differences in endothelial cell proliferation however endothelial cell nuclei were frequently stacked, pointing to altered endothelial cell distribution. Possibly, the combination of reduced tip cells guiding the angiogenic front and the continuous proliferation from stalk cells results in endothelial cell crowding. Further, no defects in generating new endothelial cells may permit *Pdgfbi*^cre/+^ *dnRAR403*^fl/fl^ mutants to ‘recover’ from early defects in vascular plexus formation, though more studies are needed to test this idea further.

Analysis of gene expression in endothelial cells showed reduced gene expression of *Vegfr3* in endothelial RA signaling mutants. Tammela and colleagues discovered that VEGFR3 is highly expressed within CNS (retinal) and non-CNS endothelial tip cells and blocking this receptor using antibodies or genetic deletion results in decreased sprouting, vascular density, and vessel branching (27), similar to what we observe in endothelial RA signaling mutants. RA via its receptor RARα has previously been shown to stimulate *Vegfr3* in lymphatic endothelial cells during lymphatic vasculature development and in lung cancer cells (28,29). Expression of Notch pathway genes and *Vegfr2* was not altered however *Lef1*, a key downstream effector of the Wnt-β-catenin pathway that has essential roles in CNS vascular development, was also decreased. This suggests that Wnt-β-catenin may be altered in endothelial RA signaling mutants however the intact BRB and no change in a different Wnt signaling readouts (*Axin2*) suggests it may play a minor role in the phenotype.

We have previously shown that during fetal brain vascular development, RA signaling has a complex interplay with the Wnt-β-catenin pathway and VEGFA pathways to regulate prenatal brain vascular development. RA acts cell-autonomously to inhibit endothelial Wnt signaling to modulate *Pdgfb* and pericyte coverage (19,30). On the other hand, RA functions non-cell autonomously by regulating expression of VEGFA by neural stem cells in the developing cortex to promote cerebral angiogenesis (31). The different actions of RA on different cell types within the CNS to regulate vascular development is further underscored by prior studies of RA signaling in retinal vascular development and BRB integrity. Using pharmacological inhibitors, several studies have identified RA inhibition impairs retinal angiogenesis, endothelial cell proliferation, and BRB integrity (17,18). These studies underscore that the angiogenic defects observed when retinoic acid is globally inhibited is potentially due to a combination of cell autonomous (angiogenesis) and non-cell autonomous (BRB). This suggests that RA is acting on other cell types to regulate the BRB such as pericytes or astrocytes. Pericytes are potential candidate since they are known to play a role in maintaining the BRB (32). Astrocytes have high RA signaling activity in the retina based on expression of the *RARE-lacZ* genetic reporter (33) and are well known to play a role in retinal angiogenesis and BRB integrity (17,33,34). Further, RA has different roles in the brain versus the retina vasculature development, however, more studies are needed to parse out how RA signaling regulates brain versus retina endothelial cells.

When looking at P60, the blood vessel area remains unchanged between controls and mutants, but there were residual collagen-4 sleeves detected in the primary plexus. Collagen-4 sleeves are important during development because they are a marker of remodeling and pruning of unwanted capillaries through selective branch regression (35). The persistence of Collagen-4 sleeves could point to decreased phagocytosis of empty sleeves by microglia or a defect in endothelial cells or pericytes to retract (35,36). More work needs to be done to determine the exact mechanism.

Our work provides important insight into the role of RA signaling in retinal vascular development and may have important implications for retinopathy of prematurity (ROP), a major cause of blindness in premature infants. ROP develops because the retinal vasculature of preterm infants, unlike term infants, is relatively immature as the peripheral retina is not yet vascularized (37). Hyperoxia therapy (85-95% oxygen) for preterm infants improves survival but interferes with hypoxia-driven expression of VEGF-A and stalls retinal angiogenesis. This leaves portions of retina without vascular support to meet the metabolic demands of the maturing retina. Upon return to normal air (21%oxygen), hypoxia in the poorly vascularized retina induces VEGF-A expression and aberrant neovascularization which, left untreated by surgery, can lead to blindness. In animal models of ROP, daily treatment with exogenous RA while exposed to hyperoxia prevents formation of neovascularization (38). Vitamin A supplementation to preterm infants, initially used to treat lung-related prematurity, decreases ROP incidence but the underlying mechanism for why it helps is not known (39). Possibly, endothelial signaling RA maintains VEGFR3 levels to regulate angiogenesis and RA (produced from Vitamin A) may also act on other cell types (astrocytes) in the retina to regulate other pathways involved in angiogenesis and BRB development. Future studies combining ROP animal models with *Pdgfbi*^cre/+^ *dnRAR403*^fl/fl^ may provide further insight into the role of RA in normal and pathological angiogenesis in the retina.

## Acknowledgements

This work is supported by funding from NIH/NINDS (R01 NS098273 to JAS), “Neuroscience Training Grant,” T32 NS099042 to CNC, Summer Research Training Program (funded by Neuroscience Technology Center at University of Colorado, Anschutz Medical Campus) for CC.

## Use of Animal Subjects

Use of animals for this study was approved by the University of Colorado IACUC (protocol #: 00034).

## References

1. Pawlikowski B, Wragge J, Siegenthaler JA. Retinoic acid signaling in vascular development. Genes N Y N 2000. 2019 Jul;57(7-8):e23287.

2. Yu PK, Balaratnasingam C, Morgan WH, Cringle SJ, McAllister IL, Yu DY. The Structural Relationship between the Microvasculature, Neurons, and Glia in the Human Retina. Investig Opthalmology Vis Sci. 2010 Jan 1;51(1):447.

3. Dai C, Xiao J, Wang C, Li W, Su G. Neurovascular abnormalities in retinopathy of prematurity and emerging therapies. J Mol Med Berl Ger. 2022 Apr 8;

4. Blanco R, Gerhardt H. VEGF and Notch in tip and stalk cell selection. Cold Spring Harb Perspect Med. 2013 Jan 1;3(1):a006569.

5. Gerhardt H, Golding M, Fruttiger M, Ruhrberg C, Lundkvist A, Abramsson A, et al. VEGF guides angiogenic sprouting utilizing endothelial tip cell filopodia. J Cell Biol. 2003 Jun 23;161(6):1163–77.

6. Iruela-Arispe ML, Davis GE. Cellular and molecular mechanisms of vascular lumen formation. Dev Cell. 2009 Feb;16(2):222–31.

7. Phng LK, Gerhardt H. Angiogenesis: a team effort coordinated by notch. Dev Cell. 2009 Feb;16(2):196–208.

8. Stefater JA, Lewkowich I, Rao S, Mariggi G, Carpenter AC, Burr AR, et al. Regulation of angiogenesis by a non-canonical Wnt-Flt1 pathway in myeloid cells. Nature. 2011 May 29;474(7352):511–5.

9. Franco CA, Jones ML, Bernabeu MO, Geudens I, Mathivet T, Rosa A, et al. Dynamic Endothelial Cell Rearrangements Drive Developmental Vessel Regression. Hogan BLM, editor. PLOS Biol. 2015 Apr 17;13(4):e1002125.

10. Hendrikx S, Coso S, Prat-Luri B, Wetterwald L, Sabine A, Franco CA, et al. Endothelial Calcineurin Signaling Restrains Metastatic Outgrowth by Regulating Bmp2. Cell Rep. 2019 Jan;26(5):1227–1241.e6.

11. Sharma S, Shen T, Chitranshi N, Gupta V, Basavarajappa D, Sarkar S, et al. Retinoid X Receptor: Cellular and Biochemical Roles of Nuclear Receptor with a Focus on Neuropathological Involvement. Mol Neurobiol. 2022 Apr;59(4):2027–50.

12. Goto S, Onishi A, Misaki K, Yonemura S, Sugita S, Ito H, et al. Neural retina-specific Aldh1a1 controls dorsal choroidal vascular development via Sox9 expression in retinal pigment epithelial cells. eLife. 2018 Apr 3;7:e32358.

13. McCaffery P, Dräger UC. Retinoic acid synthesis in the developing retina. Adv Exp Med Biol. 1993;328:181–90.

14. McCaffery P, Tempst P, Lara G, Dräger UC. Aldehyde dehydrogenase is a positional marker in the retina. Dev Camb Engl. 1991 Jul;112(3):693–702.

15. Cvekl A, Wang WL. Retinoic acid signaling in mammalian eye development. Exp Eye Res. 2009 Sep;89(3):280–91.

16. Duester G. Towards a Better Vision of Retinoic Acid Signaling during Eye Development. Cells. 2022 Jan 19;11(3):322.

17. Engelbrecht E, Metzler MA, Sandell LL. Retinoid signaling regulates angiogenesis and blood-retinal barrier integrity in neonatal mouse retina. Microcirc N Y N 1994. 2022 Apr;29(3):e12752.

18. Pollock LM, Xie J, Bell BA, Anand-Apte B. Retinoic acid signaling is essential for maintenance of the blood-retinal barrier. FASEB J Off Publ Fed Am Soc Exp Biol. 2018 Oct;32(10):5674–84.

19. Bonney S, Harrison-Uy S, Mishra S, MacPherson AM, Choe Y, Li D, et al. Diverse Functions of Retinoic Acid in Brain Vascular Development. J Neurosci Off J Soc Neurosci. 2016 Jul 20;36(29):7786–801.

20. Claxton S, Kostourou V, Jadeja S, Chambon P, Hodivala-Dilke K, Fruttiger M. Efficient, inducible Cre-recombinase activation in vascular endothelium. Genes N Y N 2000. 2008 Feb;46(2):74–80.

21. Rosselot C, Spraggon L, Chia I, Batourina E, Riccio P, Lu B, et al. Non-cell-autonomous retinoid signaling is crucial for renal development. Dev Camb Engl. 2010 Jan;137(2):283–92.

22. Zudaire E, Gambardella L, Kurcz C, Vermeren S. A Computational Tool for Quantitative Analysis of Vascular Networks. Ruhrberg C, editor. PLoS ONE. 2011 Nov 16;6(11):e27385.

23. Lee NJ, Ha SK, Sati P, Absinta M, Luciano NJ, Lefeuvre JA, et al. Spatiotemporal distribution of fibrinogen in marmoset and human inflammatory demyelination. Brain. 2018 Jun 1;141(6):1637–49.

24. MacCormick IJC, Barrera V, Beare NAV, Czanner G, Potchen M, Kampondeni S, et al. How Does Blood-Retinal Barrier Breakdown Relate to Death and Disability in Pediatric Cerebral Malaria? J Infect Dis. 2022 Mar 15;225(6):1070–80.

25. Brash JT, Bolton RL, Rashbrook VS, Denti L, Kubota Y, Ruhrberg C. Tamoxifen-Activated CreERT Impairs Retinal Angiogenesis Independently of Gene Deletion. Circ Res. 2020 Aug 28;127(6):849–50.

26. Mettouchi A. The role of extracellular matrix in vascular branching morphogenesis. Cell Adhes Migr. 2012 Dec;6(6):528–34.

27. Tammela T, Zarkada G, Wallgard E, Murtomäki A, Suchting S, Wirzenius M, et al. Blocking VEGFR-3 suppresses angiogenic sprouting and vascular network formation. Nature. 2008 Jul;454(7204):656–60.

28. Marino D, Dabouras V, Brändli AW, Detmar M. A role for all-trans-retinoic acid in the early steps of lymphatic vasculature development. J Vasc Res. 2011;48(3):236–51.

29. Kalitin NN, Karamysheva AF. RARα mediates all-trans-retinoic acid-induced VEGF-C, VEGF-D, and VEGFR3 expression in lung cancer cells. Cell Biol Int. 2016 Apr;40(4):456–64.

30. Bonney S, Dennison BJC, Wendlandt M, Siegenthaler JA. Retinoic Acid Regulates Endothelial β-catenin Expression and Pericyte Numbers in the Developing Brain Vasculature. Front Cell Neurosci. 2018;12:476.

31. Mishra S, Choe Y, Pleasure SJ, Siegenthaler JA. Cerebrovascular defects in Foxc1 mutants correlate with aberrant WNT and VEGF-A pathways downstream of retinoic acid from the meninges. Dev Biol. 2016 Dec 1;420(1):148–65.

32. Park DY, Lee J, Kim J, Kim K, Hong S, Han S, et al. Plastic roles of pericytes in the blood-retinal barrier. Nat Commun. 2017 May 16;8:15296.

33. Yao H, Wang T, Deng J, Liu D, Li X, Deng J. The development of blood-retinal barrier during the interaction of astrocytes with vascular wall cells. Neural Regen Res. 2014 May 15;9(10):1047–54.

34. Perelli RM, O’Sullivan ML, Zarnick S, Kay JN. Environmental oxygen regulates astrocyte proliferation to guide angiogenesis during retinal development. Dev Camb Engl. 2021 May 1;148(9):dev199418.

35. Simonavicius N, Ashenden M, van Weverwijk A, Lax S, Huso DL, Buckley CD, et al. Pericytes promote selective vessel regression to regulate vascular patterning. Blood. 2012 Aug 16;120(7):1516–27.

36. Rymo SF, Gerhardt H, Wolfhagen Sand F, Lang R, Uv A, Betsholtz C. A two-way communication between microglial cells and angiogenic sprouts regulates angiogenesis in aortic ring cultures. PloS One. 2011 Jan 10;6(1):e15846.

37. Hellström A, Smith LEH, Dammann O. Retinopathy of prematurity. Lancet Lond Engl. 2013 Oct 26;382(9902):1445–57.

38. Wang L, Shi P, Xu Z, Li J, Xie Y, Mitton K, et al. Up-regulation of VEGF by retinoic acid during hyperoxia prevents retinal neovascularization and retinopathy. Invest Ophthalmol Vis Sci. 2014 May 27;55(7):4276–87.

39. Darlow BA, Graham PJ. Vitamin A supplementation to prevent mortality and short and long-term morbidity in very low birthweight infants. Cochrane Database Syst Rev. 2007 Oct 17;(4):CD000501.

